# Breath to genome: Whole genome sequencing of large whales from blow sampling

**DOI:** 10.64898/2026.03.08.710374

**Authors:** Éadin N. O’Mahony, Morgan L. McCarthy, Eric M. Keen, Janie Wray, Christie J. McMillan, Sheila J. Thornton, Luke E. Rendell, Morten Tange Olsen, Oscar E. Gaggiotti

**Affiliations:** Sea Mammal Research Unit, Scottish Oceans Institute, University of St Andrews, East Sands, St Andrews, KY16 9LB, UK; Centre for Biological Diversity, University of St Andrews, St Andrews KY16 9TH, UK; Section for Molecular Ecology & Evolution, Globe Institute, University of Copenhagen, Copenhagen, Denmark; North Coast Cetacean Society, 26 Cottonwood Road, Alert Bay, BC, V0N 1A0, Canada; Sewanee: The University of the South, 735 University Avenue, Sewanee TN 37383, USA; Pacific Biological Station, Fisheries and Oceans Canada, Nanaimo, British Columbia, Canada; Pacific Science Enterprise Centre, Fisheries and Oceans Canada, West Vancouver, British Columbia, Canada

**Keywords:** Blow sampling, baleen whales, *Megaptera novaeangliae*, shotgun sequencing, low coverage whole genomes, SNPs

## Abstract

Preserving biodiversity requires the genetic monitoring of wild populations, but traditional invasive sampling techniques can impact animal welfare. For cetaceans, a promising non-invasive approach is the collection of exhaled respiratory vapour, or ‘blow’, to generate individual genetic profiles. Here, we demonstrate the feasibility of generating whole genomes from the blow of humpback whales with data of sufficient quality for population genomics. A total of 58 blow samples from 26 Northeast Pacific humpback whales (*Megaptera novaeangliae*) were collected in Gitga’at First Nation territory using a commercially available drone. The high endogenous content at 84% on average allowed us to generate low-coverage nuclear genomes (mean 2.3×, range 0.2×-3.5×) and high coverage mitochondrial genomes (mean 94×, range 7.7×-151.3×) for inference of population structure, diversity and phylogenomics. The reliability of the blow-derived genomes was demonstrated by direct comparison between replicate blow samples, as well as paired tissue samples from a subset of individuals.

## INTRODUCTION

Modern genomics has become an important tool in conservation biology with the ability to accurately profile genetic population structure, measure relatedness, and estimate effective population sizes (Cammen et al., 2016). The best genomic data are obtained from invasive tissue sampling, but for species of perilous conservation status such sampling can be undesirable and restricted for management and ethical concerns. In such cases non-invasive methods of collecting genetic samples are preferred, to avoid harm and minimise disturbance to individuals, an increasingly pertinent consideration in an era of persistent, pervasive anthropogenic pressures on nature (Byrne et al., 2021; Zemanova, 2019). Often non-invasive methods allow sampling when and/or where more conventional sampling methods (e.g. biopsies) are not possible, such as in species that are elusive or protected from harm or take (e.g. paw print sampling of polar bears *Ursus maritimus*; Von Duyke et al., 2023). Legal permitting or physical access to the habitat, such as in marine ecosystems, can also be challenging (Visser et al., 2021). Fortunately, minimally- and non-invasive genetic sampling and their ethical benefits are becoming increasingly available thanks to new sampling technologies and the rapid advances and decreasing costs of genome sequencing technologies (Carroll et al., 2018; Waits & Paetkau, 2005).

Minimally- or non-invasive genetic sampling of marine species varies between taxonomic groups of interest, with environmental (e)DNA and sedimentary ancient (seda)DNA often providing a broad-scale overview of entire communities (e.g. Port et al., 2016; Schreiber et al., 2024), while more species-specific approaches range from mucus swabs in elasmobranchs (Kashiwagi et al., 2015; Lieber et al., 2013) to fluke-footprint eDNA, sloughed skin and exhaled respiratory vapour or ‘blow’ sampling of cetaceans (whales, dolphins and porpoises) using collection poles or drones (Amos et al., 1992; Frère et al., 2010). In the context of blow sampling, drones have been applied at various latitudes to a range of cetacean species, including mysticetes – humpback whales *Megaptera novaeangliae*, fin whales *Balaenoptera physalus*, blue whales *B. musculus*, gray whales *Eschrichtius robustus* – and odontocetes – sperm whales *Physeter macrocephalus* and bottlenose dolphins *Tursiops truncatus* (Acevedo-Whitehouse et al., 2010; Atkinson et al., 2021; Centelleghe et al., 2021; Costa et al., 2022; O’Mahony et al., 2024). The blow, whether collected with a drone or an extended pole, has been used to assess whale health through the characterisation of respiratory microbiota (e.g., Acevedo-Whitehouse et al., 2010) and hormonal profiles (e.g., Burgess et al., 2018; Hogg et al., 2009). With varying levels of success, host DNA has been extracted from the blow of a range of species, all of which amplify short fragments of the mitochondrial genome and/or microsatellite loci from the nuclear genome (Atkinson et al., 2021; Borowska et al., 2014; Frère, Krzyszczyk, Patterson, Hunter, et al., 2010; Neveceralova et al., 2025; O’Mahony et al., 2024; Richard et al., 2017; Robinson & Nuuttila, 2020).

For cetacean blow to be a useful non-invasive sample source for conservation genomics, a much higher number of molecular markers than previously achieved will be necessary (Cammen et al., 2016). The development of high-throughput sequencing (HTS) has seen a shift from working with a single or small number of molecular markers, such as mitochondrial control region haplotypes and microsatellites, to whole genome analyses (Allendorf et al., 2010). Additionally, these rapidly advancing molecular techniques allow for progressively lower quantities of sample DNA to generate whole genomes (de Flamingh et al., 2022). The analysis of genomes at low depth of coverage, through the use of genotype likelihoods (Lou et al., 2021), can result in valuable population genomic insights, such as population differentiation, allele frequency estimation, relatedness, genetic diversity and inbreeding (Fuentes-Pardo & Ruzzante, 2017).

Here, we demonstrate for the first time the feasibility of obtaining low-coverage nuclear genomes and high coverage mitochondrial genomes (hereafter ‘mitogenomes’) from the blow of free ranging baleen whales to generate data of sufficient quality for population genomic inference. Samples were collected from humpback whales in Gitga’at First Nation territory in what is now known as British Columbia (BC), Canada, as described by our previous work detailed in O’Mahony et al. (2024), using a small commercially available unoccupied aerial vehicle (UAV), or drone. This method significantly reduces the disturbance to the target animals relative to current methods by removing the need for close vessel-approaches and invasive biopsy sampling (Christiansen et al., 2020).

### BOX 1. Positionality and reflexivity statement

We are an international group of marine mammal researchers united by the motivation to simultaneously minimise impact on the cetacean populations we study, while maximising the research output possible from the data we collect. We have an almost equal gender balance, and are collaborating across community-, NGO-, governmental- and academic-rooted organisations. Authors ÉOM, MLM, MTO, LER and OEG are based at European universities, whilst EMK is based at a university in the United States. Authors JW, CJM and SJT are based in Canada. All authors are grateful for our ongoing collaboration with multiple First Nation communities, particularly the Gitga’at First Nation, and extend our gratitude for the opportunity to study the recovering populations of baleen whales foraging in the fjords of Gitga’at territory in the eastern North Pacific.

## MATERIALS AND METHODS

### Approach to methodological validation

Demonstrating the reliability of blow samples to generate whole genome sequencing (WGS) data requires several analytical steps. The most important one is demonstrating that duplicate samples collected separately from the same field-identified individual are genomic replicates of each other. But it is equally important to evaluate sequencing depth and endogenous content of samples for both nuclear and mitochondrial genomes, to establish if the genetically-assigned sex of individuals matches field observations, and to compare heterozygosity and runs of homozygosity (ROH) estimates from blow samples with those calculated from biopsy samples. In addition, the low-coverage WGS (lcWGS) from the blow of two individual humpback whales were compared to medium coverage WGS from biopsy samples visually matched to the same individual (15-16× depth of coverage).

### Sample collection

Blow samples were collected in 2022 and 2023 using a small quadcopter drone (DJI Mavic 2 Pro) in Gitga’at First Nation territory, northern BC, Canada, by author ÉOM, a Transport Canada Advanced Operations drone pilot (under Department of Fisheries and Oceans research permit #XMMS12022). Blow sampling was reviewed and approved by the University of St Andrews’ Animal Welfare and Ethics Review Body (Reference #BL16194). The lid and base of a sterile petri dish was secured to custom 3D-printed drone attachments (created by Ocean Alliance, www.whale.org) shortly before each flight (Fig. 1C). With the sample collected (Fig. 1D), the petri dish was sealed using parafilm, labelled, placed in a Ziplock and stored on ice until later processing. Samples were collected opportunistically in good weather conditions (Beaufort < 3). Paired fluke-identification (ID) photographs were captured using standard methods (described in O’Mahony et al., 2024) and behavioural responses of the whale to the drone were monitored in real-time. A response scale adapted from (Weinrich et al., 1992) which we have used for blow sampling since 2019 (O’Mahony et al., 2024), was also applied here to grade levels of behavioural response of the target whales to the drone and/or our presence in the research vessel. To minimise disturbance, resting/sleeping whales were not sampled. Photo-ID was also captured in 4K video from the drone when possible and used to corroborate DSLR-collected identifications. Individual whales were typically only sampled once per encounter. Duplicate and triplicate samples were collected from individuals during different encounters, to allow assessment of sample contamination and reliability of genomic data across different group sizes and behavioural states.

**Figure 1.**
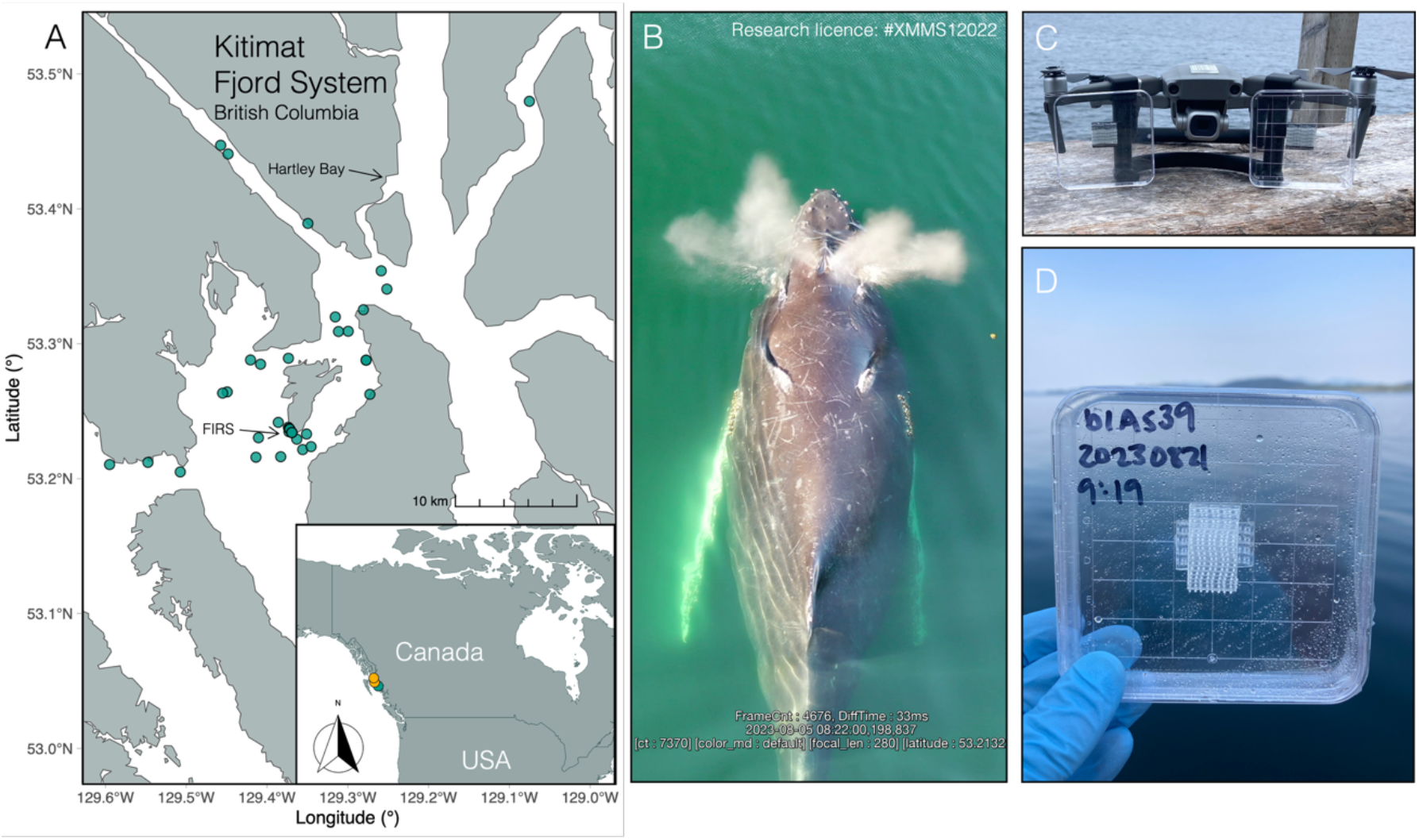
**A)** The blow sampling study area, which falls within Gitga’at First Nation territory, British Columbia, Canada. Blow sampling locations are plotted in green and biopsy sampling locations in yellow. The research station is marked as ‘FIRS’ (Fin Island Research Station) on the southern end of Fin Island. **B)** A frame extracted from a blow sampling flight to demonstrate the feasibility of using scarring visible on the back of the whale to cross-reference the individual ID between replicate sampling events. **C)** The drone with custom 3D-printed attachments to hold the sterile petri dish secure shortly before each flight. **D)** An example of humpback whale blow samples shortly after collection, before processing.

Immediately upon sample collection, the drone was returned to the operator and petri dishes sealed and placed on ice until return to the field station. Petri dishes were then rinsed with 0.5 mL Longmire’s solution and 1.5 mL distilled H_2_O using a sterile scraper to ensure all of the blow droplets were washed into solution. This solution was then pipetted into a 2 mL cryovial and frozen in a household freezer set to -20°C, until being shipped on dry ice to the laboratories of the Globe Institute, University of Copenhagen, Denmark.

Biopsy samples were collected in 2022 by the Department of Fisheries and Oceans Canada under Marine Mammal License Number MML-01. Individual whales were identified prior to sampling using standard fluke-ID photographs and techniques. Samples were collected using a hollow-tipped arrow (40 mm length, 8 mm diameter), deployed by a Barnett recurve crossbow with a draw weight of 150 lbs. The biopsy tips were cleaned and sterilized between each use following protocols approved by the Pacific Region Animal Care Committee. Both samples were placed in a cooler on ice immediately after being obtained and transferred to liquid nitrogen at the end of the day. Upon return from the field, samples were stored in a -80°C freezer until further processing.

### DNA extraction to sequencing

Samples were extracted using a slightly modified Qiagen DNeasy Blood and Tissue extraction protocol (see Supplementary Materials and O’Mahony et al., 2024). The DNA concentration of the samples was quantified with the Qubit High-Sensitivity assay. Samples were extracted with negative controls (extraction blanks) and two positive controls – both ringed seals *Pusa hispida*, sampled in Pond Inlet and Resolute, Nunavut, Canada. Humpback whale tissue samples were extracted using a phenol–chloroform extraction with proteinase K digestion (see Supplementary Materials for further detail).

Library preparation and shotgun sequencing of blow samples was performed by BGI China using 20 uL of sample extracts. Sample extracts were prepared for sequencing using the KAPA HyperPrep library preparation kit for low input DNA and shotgun sequenced (paired-end 100 bp) on the DNBSEQ platform producing 10 Gb of data per sample. Biopsies were shotgun sequenced by Genome Quebec using a PCR-free protocol on a NovaSeq with paired end 150bp reads to a target depth of 15-20×.

### Read processing and alignment

Raw read data of 58 humpback whale blow samples and two biopsy-collected samples were quality controlled using FastQC (Andrews, 2010) and then cleaned using fastp v.0.23.4 (Chen et al., 2018) with the default settings, before being mapped to the humpback whale chromosome-level nuclear genome (sequenced from a high quality tissue sample taken from a calf on the Hawaiian breeding ground of this population’s migratory corridor; Carminati et al., 2024) and separately mapped to the mitochondrial genome of the same individual (GenBank assembly GCA_041834305.1; Carminati et al., 2024) using BWA v.0.7.17 with the MEM algorithm (Li, 2013). Alignments were sorted, converted into BAM format, and exact duplicate single-end reads (usually PCR duplicates) were removed using SAMtools v.1.17 (rmdup -S) (Li et al., 2009). The endogenous content of humpback whale DNA within the blow samples was calculated as the fraction of reads mapping to the humpback whale reference genome to the total number of reads per sample. The depth of coverage (as the mean number of reads mapping to each position along the complete genome) were calculated using SAMtools (Li et al., 2009). A bed file of genomic interspersed repeats (i.e. regions of highly repetitive DNA sequences) within the humpback whale genome was created using RepeatMasker v.4.1.5 (Smit et al., 2013) and then reads mapping to these regions were masked from the cleaned bam files using BEDTools v.2.31.0 (Quinlan & Hall, 2010). We serially downsampled biopsy genomes to evaluate the effect of depth of coverage on heterozygosity and ROH using seqtk v.1.4 (Li, 2017; for further detail see Supplementary Materials).

Publicly available humpback whale genomes (n=9) were mapped to the same reference genome as the data analysed in this study. These included North (N.) Atlantic samples from a biopsy in the western N. Atlantic (8× depth; accession SRR8386009; Tollis et al., 2019), a sample from cell culture (19× depth; accession SRR5665639; Árnason et al., 2018) and six unrelated individuals from biopsy (26× depth on average; accessions SRR25114440-25114446; Suárez-Menéndez et al., 2023) and from the N. Pacific, a sample from DNA Zoo (62× depth; accession SRR17854487) and the two biopsy samples presented in this study. Raw read data was processed with the same steps as used for the blow sample dataset.

### Quality control

Several quality control (QC) steps were taken to filter first the blow sample dataset with the biopsy-sourced data and later the merged dataset (where merged samples were created from replicate blow samples).

#### Unmerged blow samples

Steps for the unmerged dataset were as follows: 1) Samples with less than 1× depth of coverage were removed. 2) Of the known replicate sets, the highest depth of coverage sample was retained. 3) Pairwise relatedness, as the KING-robust kinship coefficient (Manichaikul et al., 2010), were calculated for all remaining samples (see details in section *Relatedness*) and of the related pairs with KING > 0.125, the sample of higher depth of coverage was retained. 4) Finally, we used the ‘perfect-sample approach’ (Orlando et al., 2013) to determine if any remaining samples had relatively elevated error rates (expanded in section *Relative error rates*). We then calculated the upper 95^th^ percentile and removed any samples falling above this threshold.

#### Merged blow samples

To create the merged dataset, we followed the following QC steps: 1) We assigned mitochondrial control region haplotypes according to haplotypes used by Baker et al. (2013). Based on the field-assigned IDs, we examined pairs of replicates samples, and for any pairs with mismatching haplotypes we re-assessed ID photographs and watched the associated drone footage to confirm if the pair was in fact a mismatch based on scarring on the back of the animal (Fig. 1B). 2) In parallel to this, we sorted estimates of pairwise relatedness (see section *Relatedness* below) according to whether they are replicates (1) or not (0). Outlier pairs falling below the 25^th^ percentile threshold had their IDs re-assessed (Fig. 2B). Any pairs with confirmed mis-assigned IDs were flagged and not used to create the merged dataset. 3) Remaining replicate sets were merged using SAMTools v.1.17 *merge* to increase the depth of coverage for individual humpback whales. 4) Pairwise relatedness was recalculated using this merged dataset to flag any highly related pairs. Of pairs with a KING > 0.125, the higher depth of coverage individual was retained. 5) Finally, the ‘perfect sample approach’ was also applied to the merged samples using the same perfect individual and outgroup. The upper 95^th^ percentile was once again used as the threshold over which samples should be excluded.

**Figure 2.**
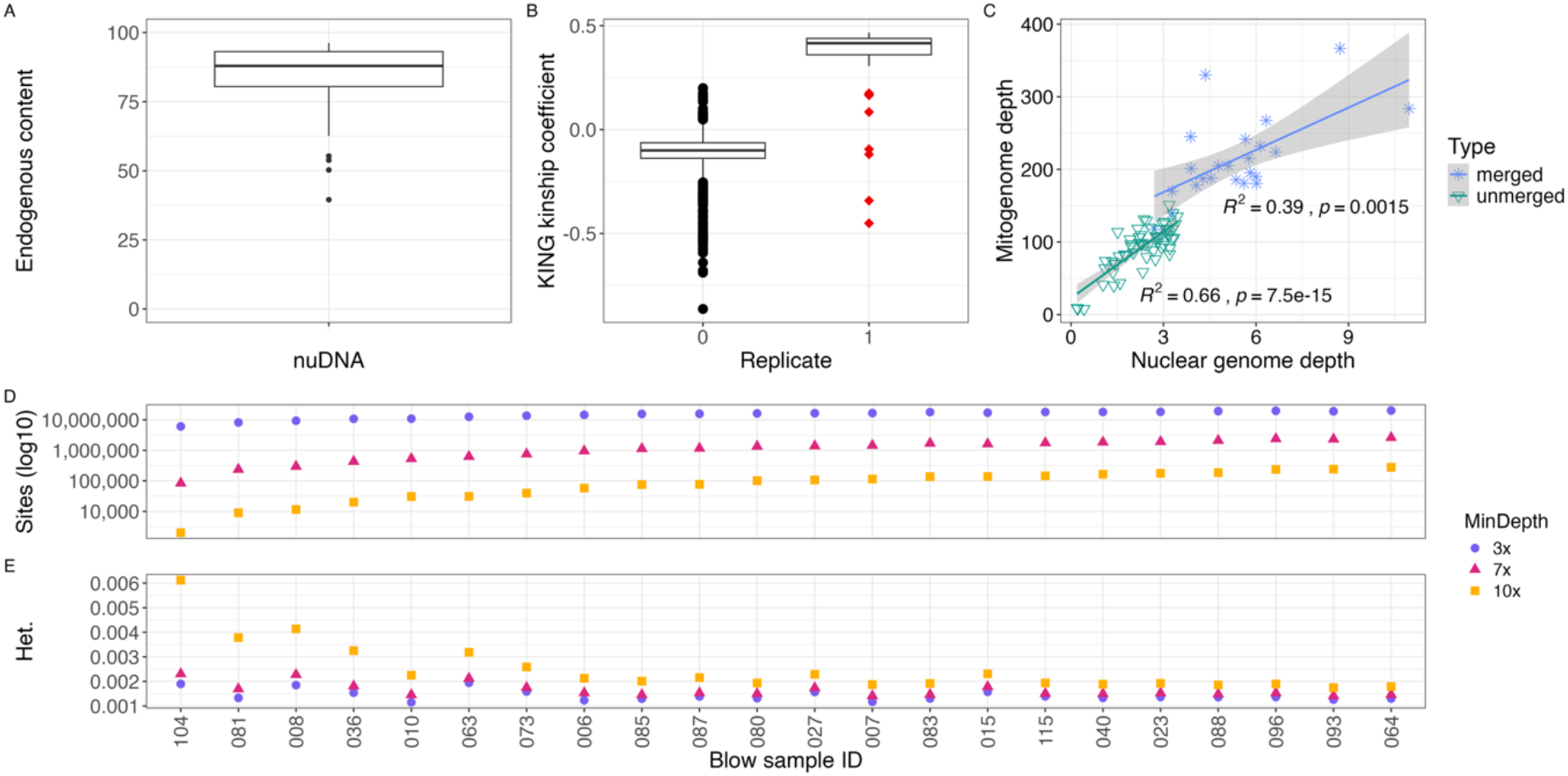
**A)** The endogenous content of unmerged blow samples for nuclear DNA (mean = 83.6% +/-12.8%). **B)** KING-robust kinship coefficients calculated without a filter for minimum depth, sorted on the x-axis according to field-assigned identification replicates (=1) and non-replicates (=0). Points highlighted as red diamonds were found to have incorrect field-assigned IDs and were removed during quality control. **C)** Relationship between nuclear and mitochondrial sequencing depth of coverage across sample types. Scatter plot showing nuclear genome depth (x-axis) versus mitogenome depth (y-axis) for each sample, coloured and shaped by dataset type (“merged” and “unmerged” blow samples). Linear regression lines are fitted separately for each type, with shaded areas representing 95% confidence intervals. Pearson’s correlation coefficient (R^2^) and associated p-values are shown for each regression. **D)** The number of sites available for autosomal heterozygosity calculation, when a filter of minimum depth of 3×, 7× and 10× was applied. **E)** Autosomal heterozygosity (Het.) of the final dataset of unmerged blow samples (worse quality replicates (i.e. lower depth of coverage) and highly related individuals removed; 22 samples remaining) using genotype likelihoods with varying per site minimum depth thresholds. Samples are ordered from left to right in an increasing number of sites available. Mean number of sites across samples: min. depth 3× = 14,979,363; 7× = 1,246,257, 10× = 101,987. Colour blind-friendly palette from R package *ggpubfigs* (Steenwyk & Rokas, 2021).

### Relative error rates

To identify samples with relatively high error rates, perhaps stemming from low depth of coverage or contamination, we used ANGSD (Korneliussen et al., 2014) to measure the distance from a ‘perfect sample’ to an outgroup and used this to quantify deviations in the frequency of derived alleles in each sample relative to the perfect sample. We used the consensus fasta of the fin whale DNA Zoo reference genome (SRR16970344; Dudchenko et al., 2017, 2018) mapped to the humpback reference as the outgroup (-anc) and the highest depth of coverage N. Pacific whole genome (SRR17854487; Dudchenko et al., 2017, 2018) as the perfect individual (-ref). We generated consensus fasta sequences using the ANGSD option -doFasta 2, which samples the most common base. For the error rate estimation, we followed standard thresholds – using a randomly selected base with -doAncError 1, a mapping and base qualities threshold of 30 and using only autosomes (-minMapQ 30 - minQ 30 -rf ${autosomes}).

### Mitochondrial genomes

A *Megaptera novaeangliae* mitogenome from the same reference genome was downloaded from NCBI (GenBank accession PP475430). Prior to mapping, a 70 bp pad was added to each end of the linearized mitogenome to ensure sufficient mappability across the artificial break of the circular mitogenome. The reference mitogenome was first indexed using SAMtools v.1.17 *faidx* (Li et al., 2009) and BWA v.0.7.17 *index* (Li, 2013) and blow sample, biopsy and publicly available raw read data were mapped using BWA *mem* (Li, 2013). Duplicates were removed (*samtools rmdup*) and once the depth (+/-one standard deviation) was calculated for each sample, a consensus mitogenome was called using ANGSD v.0.940-2 (-doFasta 2, calling the most common base at each site across all reads; Korneliussen et al., 2014).

Consensus mitogenomes were imported into Geneious Prime v.2025.0.3 and a multiple alignment was performed using the MAFFT v.7.490 alignment algorithm with default settings (Katoh et al., 2002; Katoh & Standley, 2013). We included publicly available humpback whale whole mitogenomes (n=10; GenBank accession numbers Table S4) and two N. Atlantic fin whale mitogenomes as an outgroup (GenBank accession numbers: NC_001321 (Valverde et al., 1994) and OZ239533, submitted by the Darwin Tree of Life Project). Mitochondrial control region haplotypes were assigned according to the haplotypes described in Baker et al. (2013) and haplotype frequencies calculated for comparison to frequencies found in the northern BC region of this study. The post-QC subset of samples, plus sample MN-b-070 (because all control region haplotypes were then represented by at least one sequence) and the publicly available mitogenomes were exported as a Phylip alignment and a maximum likelihood phylogenetic tree constructed using IQ-TREE (Nguyen et al., 2015) with branch support calculated using 100 replicate bootstraps. Figures 3B and S6 were created using FigTree v.1.4.4 (http://tree.bio.ed.ac.uk/software/figtree/).

**Figure 3.**
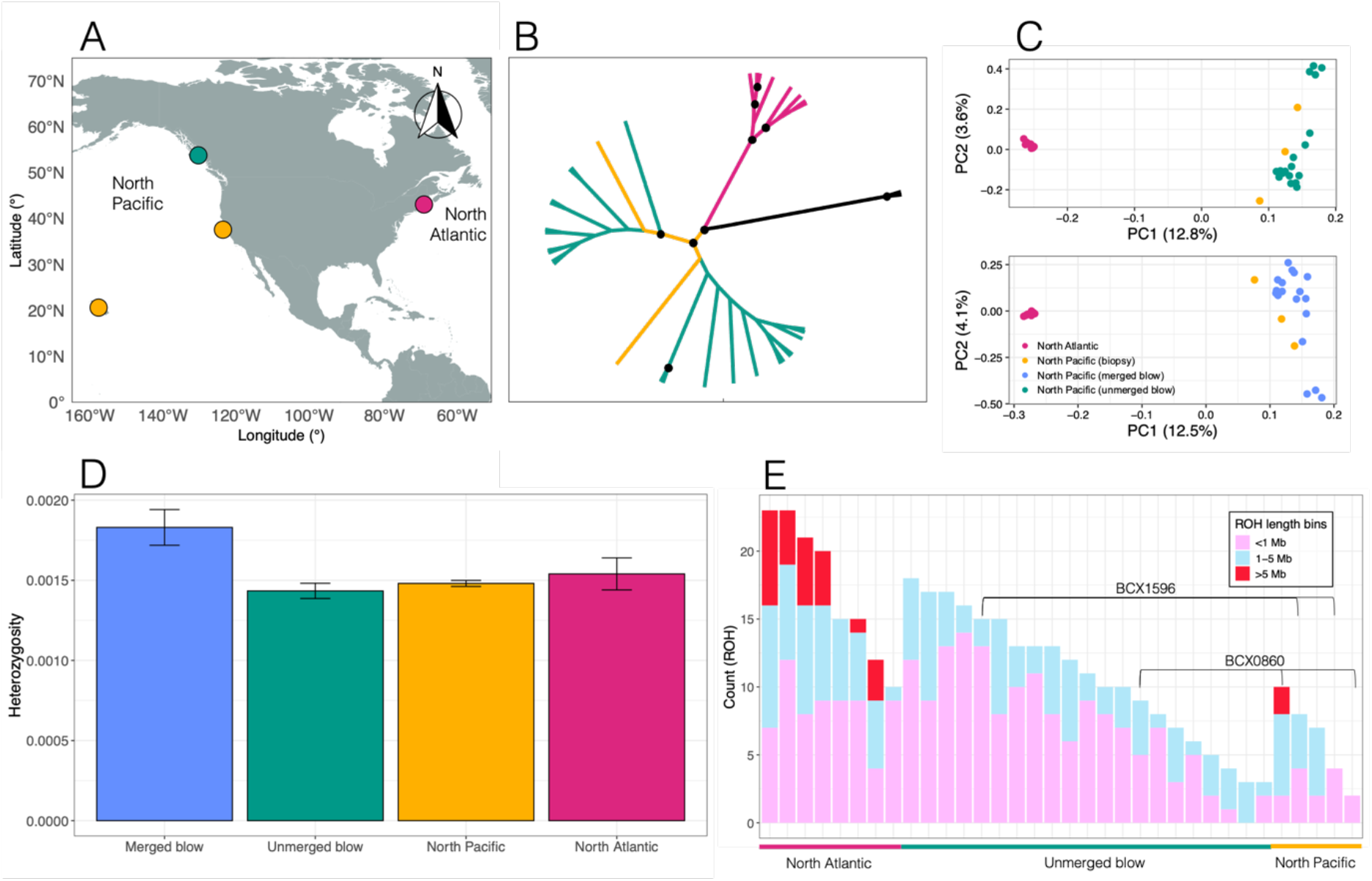
Blow samples for population genomics and phylogenetics. **A)** Map of North America showing sampling locations for blow samples collected in British Columbia, Canada (green), for the two publicly available whole genomes from the North Pacific (yellow) and for the publicly available data from the North Atlantic (pink). **B)** A cladogram constructed from consensus mitochondrial genomes, coloured according to panel A. Two North Atlantic fin whales are included as an outgroup and shown in black. Black circles indicate nodes with bootstrap support greater than 95% across 100 replicates. **C)** A principal component analysis (PCA) using both the unmerged blow samples (green) and beneath this the merged blow samples (blue), as well as novel and publicly available genomic data based on biopsy samples from the North Pacific (yellow) and the North Atlantic (pink; see Table S4 for data sources). The PCAs were generated using genotype likelihoods at 5,936,486 SNPs and 6,348,584 SNPs for the unmerged and merged blow samples, respectively. **D)** Autosomal heterozygosity of humpback whales estimated from blow samples (merged in blue and unmerged in green) compared with those generated by traditional biopsy methods. **E)** Counts of the runs of homozygosity (ROH) per individual humpback whale (unmerged blow samples and biopsy samples), coloured according to their respective lengths (pale pink < 1 Mb; pale blue 1-5 Mb and red > 5 Mb). Samples are clustered according to their respective source populations/sample type. For comparison we have highlighted the blow samples with their associated biopsies, including both the subsampled 2× biopsies and full coverage biopsies for each whale (BCX0860 and BCX1596). The blow samples and 2× subsamples have higher counts of ROH than their respective higher depth biopsy.

### Relatedness

We estimated pairwise relatedness amongst all possible pairs of blow samples and biopsies (including biopsies subsampled to lower depths of coverage) using no minimum depth threshold, with the goal of comparing relatedness coefficients between pairs of field replicates (i.e. same assigned field ID, even if sampled on different encounters) with those between pairs of non-replicate samples. This demonstrates the reliability of blow-sourced genomes, given that the sequenced genomes sampled from the same individual during different field encounters remains consistent. Estimation of relatedness also formed a QC step, as above we were able to identify incorrect IDs made in the field by lower than expected pairwise relatedness coefficients between blow samples that were identified as replicates.

We used NgsRelate v2.0 (Korneliussen & Moltke, 2015) to calculate KING-robust kinship coefficients (Manichaikul et al., 2010) to identify both replicate samples and related individuals. Initially ANGSD is used to estimate genotype likelihoods (-doGlf 3) which are then leveraged by NgsRelate in a maximum likelihood framework to calculate R0, R1 and KING coefficients. Through the use of genotype likelihoods, NgsRelate takes the uncertainty of the genotypes into account, which is particularly important when working with low-coverage data, as inferring genotypes can be highly error prone and lead to an underestimation of heterozygous sites (Nielsen et al., 2011).

Pairs of samples with a KING-robust kinship coefficient > 0.125 were examined and the lower depth sample was discarded before further analysis. This included discarding the lower quality sample of any replicate pairs. For the later analyses using merged replicate blow samples, outliers were identified using the standard Tukey method applied within each replicate group. Specifically, for each level of the replicate factor, the first (Q1) and third (Q3) quartiles of the KING coefficient distribution were computed, and the interquartile range (IQR) was defined as IQR = Q3 - Q1. A data point was classified as an outlier if it fell below Q1 - 1.5 × IQR or above Q3 + 1.5 × IQR.

The eight replicate outlier pairs were then re-examined to see if a false identification had been made in the field. This included the careful assessment of drone-collected body imagery, because the scarring on the back and rostrum of the individual can be used to corroborate a correct matching ID (Fig. 1B). For visualisation, replicate pairs were colour-coded as red diamonds if a false field identification was confirmed (Fig. 2B). These red outliers were removed before a non-parametric Mann-Whitney U test with continuity correction was used to test the null hypothesis that replicate pairs have no higher relatedness that non-replicates, which would invalidate the method, with our expectation being that replicate pairs will have a significantly higher KING-robust kinship mean than non-replicate pairs. If a false ID was confirmed, the pair was no longer considered a replicate set and therefore not included in the merged dataset. For all remaining replicate pairs, the blow sample mapped bam files were merged using SAMtools v.1.17 *merge* (Li et al., 2009).

### Sex assignment

To estimate the sex of sampled individuals, we compared normalized read depths between the X chromosome and an autosome of similar length. Chromosome sizes in the *Megaptera novaeangliae* reference genome were manually inspected, and chromosome 6 was selected as the closest in size to the X chromosome (chromosome 6 is 89.4% the size of chromosome X). We used SAMtools v.1.17 *idxstats* to extract per-chromosome read counts from aligned BAM files. The ratio of reads mapping to the X chromosome versus chromosome 6 was then computed for each individual. Based on expected genomic configurations, females (XX) were anticipated to exhibit a ratio near 1, and males (XY) a ratio near 0.5. We compared previously determined sexes (from O’Mahony et al. (2024) and known mothers) to those assigned here and computed a given error rate from the discrepancies found.

### Heterozygosity

We used realSFS implemented within ANGSD v. 0.940-2 (Korneliussen et al., 2014), using the default settings, to estimate the site frequency spectrum (SFS). This was done using saf files generated by ANGSD and using strict quality thresholds (-GL 2 -doSaf 1 -minMapQ 30 -minQ 30 -skipTriallelic 1 -uniqueOnly 1 -remove_bads 1 -only_proper_pairs 1) and for three different thresholds of minimum read depth at each considered site (minimum depth of 3×, 7× and 10×; - setMinDepth). We used the GATK genotype likelihood model (-GL 2) for SFS-based inference because it has been shown to outperform the SAMtools model when the average coverage of samples is low (≤4×; Lou et al., 2021). We used the per-sample proportion of heterozygous loci from this one-dimensional SFS to estimate heterozygosity for each individual, following methods in Pečnerová et al. (2024). To quantify the impact of sequencing depth on heterozygosity estimates, we serially subsampled our two mid-coverage biopsy genomes (15-16×) to lower depths (0.25×–15×, at 0.25×, 0.5×, and 1× intervals), calculated per-site heterozygosity at each depth, and fit a linear regression model with sequencing depth as a continuous predictor.

To place the estimated heterozygosity of blow samples in the context of publicly available data, raw reads of available humpback whale genomes were pulled from the NCBI database (Table S4). The data was mapped to the same reference genome as the blow samples and filtered using the same pipelines. There was only one additional whole genome available stemming from the N. Pacific (DNA Zoo, accession number SRR17854487).

### Principal component analysis

We ran a principal component analysis (PCA) on the post-QC individuals using PCangsd v.1.36.3 (Meisner & Albrechtsen, 2018) with genotype likelihoods (thereby accounting for missingness due to the low coverage of the blow samples). We did this first for the unmerged dataset (n=22) and again for the merged dataset (n=20), to determine expected population clustering of blow samples from the N. Pacific separate from the N. Atlantic (Fig. 3C). For these analyses, we added the mid-coverage nuclear genomes from our two biopsied animals (and omitted their blow genomes), as well as publicly available mid-to high-coverage nuclear genome data from N. Pacific and N. Atlantic humpback whales (see Table S4 for accession numbers). These analyses had second-degree relatives removed (KING coefficient > 0.125; discussed in section *Relatedness*). We did not expect to see any population structuring evident amongst the N. Pacific individuals due to the assumption that all individuals stem from one interbreeding population and have been sampled within a relatively small geographical area, in comparison to the range of the N. Pacific-wide population.

We used ANGSD v.0.940-2 (Korneliussen et al., 2014) to generate genotype likelihoods using the GATK model (-GL 2) in beagle format (-doGlf 2). Strict quality filters were applied as described in section *Genotype likelihood estimation*, with additional filters including a low minor allele frequency (MAF) threshold of 5% and a likelihood ratio test for SNPs, retaining only sites where the probability of being polymorphic is high, i.e. a p-value ≤ 1e-6 (-minMaf 0.05 -SNP_pval 1e-6). The minimum depth threshold was set to 3× (-set MinDepthInd 3) for both unmerged and merged runs.

### Runs of homozygosity

To determine the efficacy of blow-derived genomes for estimation of inbreeding, we used ROHan (Renaud et al., 2019) to estimate the ROH present in the unmerged blow, biopsy and public genomes. We calculated the proportion of the genome in ROH and the mean ROH length per individual to assess patterns of homozygosity and potential inbreeding. We further counted the ROH present in each sample and categorised them according to their length (< 1 Mb, 1-5 Mb and > 5 Mb). For comparison between low and medium coverage genomes, we additionally estimated ROH in the biopsy samples which had been subsampled down to 2× and highlighted these in our summary figure (Fig. 3E). This includes the whale BCX0860 from blow and biopsy at 2× and 15×, and whale BCX1596 from blow and biopsy at 2× and 16×. We used R v.4.4.0 (R Core Team, 2015) and ggplot2 (Wickham, 2016) for all visualisations.

## RESULTS

### Sample collection and processing

We collected 58 blow samples from 26 unique field-identified humpback whales between July and September of 2022 and 2023 in the Kitimat Fjord System of northern BC, Canada, through research agreements between the Gitga’at First Nation and the North Coast Cetacean Society (Fig. 1A-B). Of these samples, 75% were collected in 2023 and the majority during the month of August (73%). Of the collected blow samples, 22 were collected during flights launched from land and 37 collected during flights launched from the bow of our research vessel (a 12 m aluminium skiff).

For each flight, the number of exhalations sampled was maximised from the same surfacing event of the same individual, with the drone pilot and research assistant maintaining visual contact of the target whale (1-5 breaths sampled per surfacing event). When sampling from a group of whales, the most upwind animal was sampled to avoid sample contamination. Average whale group size during sampling events was 1.7 (min=1; max=5). Real-time behavioural responses were monitored by the drone pilot, graded according to the same scale used in O’Mahony et al. (2024) and the flight was terminated if a response was detected.

### Sequencing depth and endogenous content

Our dataset (n=58) included replicate blow samples collected from the same field-identified individuals (25 individuals with replicate samples in total). Across the blow sample dataset, we got an average of 55.6 million read pairs, corresponding to 111.2 million total reads, yielding an average endogenous content (i.e. percentage of reads mapping to the host species) of 83.6% (std. dev.=12.8%; min=39.5%; max=96.2%, Fig. 2A). This left an average of 18,090,163 reads (16.4% of total reads) stemming from the microbiome of the respiratory tract of the sampled whale and from residue seawater, not analysed here.

The humpback whale low-coverage nuclear genomes obtained from the 58 blow samples had a mean depth of 2.3× (± 2.4× std. dev; min=0.2×; max=3.5×; see Table S1). After the QC removal of three samples of less than 1× coverage, the blow samples had a mean depth of 2.45× (± 2.3× std. dev; min=1.03×; max=3.5×). There was a highly significant positive correlation between a sample’s depth of coverage and its endogenous content (Spearman’s rank correlation rho=0.7, *p* < 10^-16^; Fig. S1). The two biopsy samples had depths of 15× and 16× respectively and were subsequently serially downsampled to a minimum of 0.25×, to evaluate how sequencing depth impacted the estimation of heterozygosity (Fig. S7). Merging of replicate blow samples from the filtered dataset resulted in genomic data with a mean depth at 5.6× and a variance of 4.2× (min=2.7×; max=11×; Table S1). For a single individual (BCX1484), we obtained a coverage of 11× based on four blow samples: two collected in 2022 and two in 2023.

### Mitogenome haplotypes and phylogenomics

When the unmerged pre-QC blow sample dataset was mapped to a humpback reference mitogenome, the mean endogenous content was 0.04% (± 0.012% std. dev.) and the mean depth across the blow samples was 94× (± 16.3× std. dev; min=7.7×; max=151.3×). We found a strong positive correlation between the depth of coverage of the nuclear genomes and the mitogenomes for the unmerged samples (R^2^ = 0.66, *p* = 7.5e-15) and merged samples (R^2^ = 0.39, *p* = 0.0015; Fig. 2C).

When the dataset was filtered to unique individuals only but retaining samples MN-b-089 and MN-b-001 (as these had an unknown IDs), three known mtDNA control region haplotypes were present (A- = 48.3%; A+ = 27.6%; E2 = 10.3%), and we also found three new haplotypes (N1 = 6.9%; N2 = 3.4%; N3 = 3.4%). De novo mutations evident as heteroplasmy characterise these three new control region haplotypes and occur at nucleotide positions 15,640, 15,802, and 15,822 respectively (Table S2). Each of the new haplotypes were only observed in single individuals but were consistent across replicate samples of those individuals. This, along with the high depth of coverage of the mitogenomes, drastically reduces the likelihood of this signal stemming from sequencing error, although further investigation of these haplotypes is warranted. Suárez-Menéndez et al. (2023) also detected three new haplotypes stemming from heteroplasmy in their study of N. Atlantic humpback whales, albeit at different nucleotide positions along the mitogenome. For comparison, Baker et al. (2013) found similar haplotype frequencies in northern BC of A- = 57.5%; A+ = 32.1%; E1 = 0.9%; E2 = 6.6%; F2 = 0.9% and total heteroplasmy = 1.9%, from 106 samples collected primarily by biopsy dart (and some sloughed skin). We found a high bootstrap support (100%) for the phylogenetic separation of the N. Atlantic from the N. Pacific humpback whales and a further subdivision of the N. Pacific into two clusters. One N. Pacific cluster contained all A+, E2, N1 and N3 mtDNA haplotypes while the other cluster contained the A-and N2 haplotypes (Fig. S5).

### Sex assignment and relatedness

Sex of the sampled individuals was assigned by comparing the ratio of reads mapping to the X chromosome (135,787,239 bp, GenBank accession number CM085716.1) with those mapping to another chromosome of similar length (121,350,076 bp), chromosome 6 (accession number CM085700.1). Females will have a ratio closer to 1 (due to having two X chromosomes) while males will have one closer to 0.5 (XY chromosomal configuration; Fig. S2). Of the whales with known sex (either assigned by blow sample in O’Mahony et al. (2024) or in the field as observations of an adult mother closely associated with a calf), only one individual (with two blow samples) had contradicting sex assignment (error rate = 8%).

To estimate pairwise relatedness, over 6.5 million sites had genotype likelihoods assigned in preparation for the NgsRelate run (sites = 6,554,590). We re-evaluated the photo-ID and drone footage of any ‘replicate’ pairs with mismatching mtDNA haplotypes and low relatedness coefficients and discarded the mismatched sample from the merged dataset (Table 1 and red diamond-shaped outliers in Fig. 2B). An overall error rate of misidentification in the field was 4 out of 56 samples (7%). Once misidentified pairs were reassigned as non-replicates, non-replicate pairs had a mean relatedness coefficient of - 0.11, while replicate pairs had a mean relatedness of 0.41 (Fig. S3), a highly statistically significant difference (Mann-Whitney U test *W*=0, *p* < 2.2e-16).

**Table 1.** The use of blow sampling generated nuclear and mitochondrial genomes to identify misidentified individuals, which is then corroborated using the 4K video footage captured by the drone during the sampling flight (also see Fig. 1B-C).

| Sample | Field-ID | Haplotype | mtDNA Depth (± std. dev) |
| --- | --- | --- | --- |
| MN-b-006 | BCX0147 | N1 | 119.7 (± 22.6) |
| MN-b-070 | | N3 | 63.9 (± 17.3) |
| MN-b-002 | BCX0427 | A+ | 139.6 (± 35.1) |
| MN-b-089 | | E2 | 58.3 (± 11.0) |
| MN-b-090 | | A+ | 127.8 (± 15.5) |
| MN-b-001 | BCY0474 | A- | 85.4 (± 36.9) |
| MN-b-084 | | A+ | 69.7 (± 14.2) |
| MN-b-085 | | A+ | 117.5 (± 15.9) |

### Genome quality control and validation

Of the 58 blow samples sequenced, three samples were excluded because they had < 1× depth of coverage, 30 samples were removed as the lower depth replicates, two samples were excluded due to belonging to highly related pairs (KING > 0.125) and a further two samples were excluded because they fell above the 95^th^ percentile relative error rate using the ‘perfect sample approach’ (Orlando et al., 2013). This left 22 unmerged blow samples for population genomic analyses (Fig. S4A).

When replicate samples were compared for mtDNA control region haplotypes, three individuals field assigned as having multiple blow samples were found to have contradicting haplotype assignments, despite high mitogenome coverage, which alerted us to potential misidentifications in the field. Two of these individuals (BCX0427 and BCY0474) still had two correctly matched blow samples which could be merged (Table 1). This was confirmed using pairwise relatedness estimates of the nuclear whole genomes, where pairs of samples with mismatching mtDNA haplotypes had lower than expected relatedness.

For the QC of the merged dataset, three pairs of individuals were highly related (KING > 0.125; Table S3); and two individuals fell above the error rate threshold and thus these were removed from further analysis (Fig. S4B). This left a total of 20 individuals in the final merged dataset. All samples flagged by the perfect sample approach also had elevated estimates of heterozygosity (Table S1).

### Population genomics

We demonstrated the applicability of our blow-derived low-coverage nuclear genome data for population genomic studies by inferring population structure and genetic diversity. We found a clear separation of clusters according to ocean-basin, suggesting no population structure amongst the samples represented from each area (Fig. 3C and S6). We estimated autosomal heterozygosity for each individual humpback whale in the unmerged and merged datasets and placed these estimates in the context of heterozygosity estimates from the biopsy data and high coverage data using publicly available humpback whale genomes (Fig. 3D). We found the merged samples to have an elevated measure of heterozygosity when compared with unmerged blow samples and biopsy samples, suggesting further incorrect field identifications and as a consequence the merging of samples stemming from different individuals (as discussed above). We therefore recommend working with unmerged blow samples to avoid this potential error.

The unmerged blow sample mean heterozygosity across samples was 0.00143, which was slightly lower than the biopsy sample estimates of 0.00148 and 0.00154 stemming from the N. Pacific and N. Atlantic respectively. Estimates of heterozygosity presented here for the blow samples had a minimum depth threshold of 3× at each site (retaining a mean number of sites for genotype likelihood estimation = 14,979,363). We chose this threshold because when we varied the minimum depth threshold, which greatly affected the number of SNPs available for analysis, we found a minimum depth threshold of 3× to generate the closest estimates of heterozygosity compared with higher coverage data (Fig. 2D). The mean number of sites across samples that were available for heterozygosity calculation at the various depth thresholds were as follows: min. depth 3× = 14,979,363; min. depth 7× = 1,246,257, and min. depth 10× = 101,987.

Heterozygosity estimates of subsampled biopsy-sourced genomes increased significantly with sequencing depth (β = 1.08 × 10^−5^ per 1× coverage across 0.25×–15×, R^2^ = 0.44, *p* = 1.6 × 10^−6^), indicating systematic underestimation at lower coverage (Fig. S7). Heterozygosity estimates plateaued at approximately 4.5×, suggesting deeper sequencing of blow samples to at least this depth of coverage would result in more reliable data for this type of analysis.

The unmerged blow samples had on average 0.7% (95% CI: 0 - 7.9%) of their genomes in ROH, with an average ROH length of 1.5 Mb. This low proportion of the genome in ROH, together with the moderate average ROH length, indicates minimal recent inbreeding and suggests that homozygosity in these individuals is more likely the result of distant shared ancestry or historical demographic events such as population bottlenecks. The two biopsy-sourced genomes (BCX0860 and BCX1596) had 0.08% and 0.16% of their genomes in ROH respectively and an average ROH length of 1.0 Mb. For comparison, the single publicly available genome from the N. Pacific (DNA Zoo, GenBank accession SRR17854487) had 0.56% (95% CI: 0.2 - 0.7%) of its genome in ROH with an average ROH length of 1.8 Mb. The publicly available data from the N. Atlantic had on average 3.7% (95% CI: 3.1 - 4.5%) of their genomes in ROH, with an average ROH length of 4.7 Mb. We found lower counts of ROH in the biopsies (at full depth of 15× and 16×) compared with the individuals’ blow samples (Fig. 3E). The counts of ROH were also elevated in the 2× biopsy subsample, suggesting higher coverage data is necessary to achieve accurate estimates of ROH and subsequent interpretation regarding inbreeding events in the population.

## DISCUSSION

We have established that a complete whale genome can be recovered from exhaled respiratory vapour (‘blow’), marking a significant methodological advance in non-invasive cetacean genomics. Prior to this, biopsies, strandings, or historical specimens were required to obtain nuclear genomic data—each with inherent limitations in availability, ethics, or sample quality (Brown et al., 1994; Clapham & Mattila, 1993; Keighley et al., 2021; Papastavrou & Ryan, 2023). This approach opens new avenues for conservation genomics, population monitoring, and real-time microbiome-based health assessments in the wild and can fill critical data gaps in cetacean genomic databases. By demonstrating the feasibility of lcWGS from blow samples collected using a drone, our study contributes towards addressing current needs for population genomics in management and conservation, the increased adoption of hologenomic data in evolutionary biology and animal health studies, as well as a valuable data-archive for future studies.

Several tests demonstrated the reliability and replicability of the blow-sourced genomic data, showing that the blow of free ranging whales can contain enough host DNA to generate on average 2.3× and up to 3.5× depth of coverage despite only a relatively low amount of data having been generated per samples (10 Gb per sample). Further validation was provided by the collection of replicate samples from the same individual humpback whales across two foraging seasons (25 individuals with 2-4 replicate samples each). We show that replicate pairs are genomic duplicates of each other with pairwise relatedness being significantly higher than those of non-replicate pairs (*p* < 2.2e-16), despite their low depth of coverage.

The endogenous content of the blow samples was 84% on average (std. dev.=12.8%; min=39.5%; max=96.2%), leaving us with an additional average of >18 million reads per sample stemming from the microbiome of each whale’s respiratory tract. This can further the study of cetacean holobionts, as from a single breath sample both host and microbiome genomes can be analysed. Paired with routine blow sampling for targeted study of hormonal profiles present within the breath (Burgess et al., 2018), blow becomes a valuable biological sample to assess individual to population-level health and resilience in the face of increasing anthropogenic pressures, including a rapidly changing global climate.

The collection of blow is increasingly seen as an alternative source of biological data compared with traditional biopsy sampling, which is particularly beneficial in contexts in which biopsy sampling is not viable (for example due to strict non-invasive research protocols or where permitting and training pose a challenge). Blow sampling of cetaceans thus far has yielded data on hormonal profiles representative of life-history stages (Burgess et al., 2018); microbiota found within the respiratory tract (Apprill et al., 2017); and DNA of high enough quality to assign sex, mitochondrial haplotypes and enough microsatellite loci for individual identification (on average 1.7 loci across multiple species (Atkinson et al., 2021) and 7.5 loci for humpback whales; O’Mahony et al., 2024). The added benefit of blow collection using a drone is that one can eliminate the need for a boat if flying from land (37% of samples in this study). Or, at minimum, avoid the need for a close vessel approach, which is required for both pole-based blow sampling and biopsy sampling (Hogg et al., 2009; Noren & Mocklin, 2011). Given the size and mass of cetaceans, a close vessel approach poses risks to both the welfare of the researchers and the target species (Pirotta et al., 2017).

Additionally, blow genomic data can be utilised for genetic mark-recapture studies (Palsbøll et al., 1997), as well as ground truthing of field photo-IDs of individual whales. Here we had a small handful of instances in which estimated relatedness and mismatching mitochondrial haplotypes alerted us to putative false IDs made in the field. Correct field IDs can be challenging to obtain in certain circumstances, such as when there is a large group of whales foraging together or if an individual whale does not fluke. The use of a drone for sampling, which allows for the simultaneous recording of the flight in 4K video, presents the opportunity to recheck field IDs of sampled whales using the distinct markings on the rostra and dorsal regions of the whales, even years after the sampling event (Fig. 1B; and see Degollada et al., 2023; O’Mahony et al., 2024).

Next-generation sequencing has revolutionised the fields of ecology and evolution in model and non-model organisms, especially the way that probabilistic genotype likelihood-based tools, such as ANGSD (Korneliussen et al., 2014) and ROHan (Renaud et al., 2019), give us the opportunity to carry out population genomics analyses on lower coverage data. Indeed, we have shown that despite sequencing the individual blow samples to relatively low coverage, a large number of SNPs are still available to carry out various population genomics tests – including PCAs which parse out the different populations, N. Atlantic and N. Pacific, as expected (Fig. 3C) and measures of heterozygosity which correspond with estimates stemming from higher coverage biopsy-sourced genomes (Fig. 3D).

In addition, we have generated a resource database of low-coverage whole genomes for N. Pacific humpback whales which can be improved upon through genotype imputation using future high coverage data – something that is not possible with microsatellite data. However, given the high endogenous content of our blow samples, higher coverage genomes could also be obtained at little additional economic costs, allowing for more in-depth population genomic analyses such as modelling of demographic histories and estimation of genetic load. We recommend obtaining single blow samples with as many exhales as possible in the same sampling flight, and sequencing these to a deeper coverage. This will also mitigate the risk of merging of samples from different individuals stemming from a false field-ID, which most likely explains the elevated heterozygosity visible in our merged blow dataset (Fig. 3D).

Our demonstration that whale blow can yield complete genomes expands the possibilities of non-invasive sampling in marine biology. This approach enables population-scale genomics, longitudinal monitoring, and evolutionary studies in cetaceans—without physical disturbance. Crucially, it offers a powerful tool for conservation, facilitating *in situ* assessments of population structure, genetic diversity, and health—key metrics for informed management in a rapidly changing ocean.

## Supporting information

O'Mahony-etal-2026-SupplementaryMaterials-preprint

## ACKNOWLEDGEMENTS

We take this opportunity to thank the Gitga’at First Nation for their ongoing stewardship of their territory in which this work was carried out and for our continued collaboration to research and protect the whales foraging within the fjords. We would like to thank the field teams at the Fin Island Research Station for their hard work and dedication to the whales of Gitga’at territory. A special thank you to Grace Baer, Nell O’Mahony, Florent Nicolas, Alma Foley, Abbey Lewis, and Jemima Beddoe for their good humour and hard work in the field. We thank Chris Zadra from Ocean Alliance for his technical assistance, 3D-print design and creation of drone attachments and contributions to field work. We would like to thank Andreas Bak Pørksen for his contributions to lab work, as well as staff at the Globe Institute Molecular Biology Labs for their continued support. The computations were carried out on the HPC cluster ‘Mjolnir’ of the Globe Institute, University of Copenhagen, and we are grateful to its maintenance and the technical support provided by Bent Petersen.

## DATA AVAILABILITY

Scripts are deposited on GitHub (https://github.com/eadinomahony/whale-blow-genomics-method) and raw sequencing data will be uploaded to the European Nucleotide Archive upon publication.

## AUTHOR CONTRIBUTIONS

Conceptualisation: ÉOM, MLM, MTO and OEG. Methodology: ÉOM, MLM, OEG and MTO. Blow sampling and associated lab work: ÉOM. Biopsy sampling: CJM and SJT. Formal analysis: ÉOM. Validation: ÉOM, MLM, LER, MTO and OEG.

Resources: ÉOM, JW, CJM, SJT, OEG and MTO. Data curation: ÉOM. Visualisation: ÉOM and MLM. Writing – original draft: ÉOM. Writing – review and editing: ÉOM, MLM, EMK, JW, CJM, SJT, LER, MTO and OEG. Supervision: MLM, LER, OEG and MTO. Project administration: ÉOM. Funding acquisition: ÉOM, JW, OEG and MTO.

The authors declare no competing interests, financial or otherwise.

