## Supplementary material for "Breath to genome: Whole genome sequencing of large whales from blow sampling": O'Mahony-etal-2026-SupplementaryMaterials-preprint

### SUPPLEMENTARY MATERIALS

#### *Additional detail on sample collection*

We aimed to collect as many breaths as possible from the same individual in the same sampling flight because O'Mahony et al. [1] previously demonstrated a positive correlation with number of breaths and sample quality. At times the behaviour of the target individual does not lend itself to collection of multiple breaths, limiting the volume of sample collected. We encountered sampling instances in which a whale would surface and exhale just twice before fluking, marking the end of the sampling window. In such cases the first breath is used to locate the whale by the drone pilot and position the drone aft of the blow hole so that the second breath can be sampled before the whale flukes. It is vital to avoid sample contamination by accidentally sampling multiple individuals, hence the sampling window ends once the whale flukes and dives out of sight.

The approach presented here comes with some challenges which should be further explored. Blow sampling of whales at sea is inherently weather dependent, and sampling with a drone gets increasingly difficult as the wind picks up. Nonetheless, this is also true for biopsy sampling, where wind, chop and swell all contribute to increasingly difficult sampling conditions. For blow sampling, the plume of exhalant gets flattened closer to the sea surface in higher winds, requiring the pilot to navigate the drone closer to water. We therefore developed a Standard Operating Procedure (SOP) which prevents blow sampling in Beaufort ≥ 3, and if sampling from a research vessel, the SOP includes a drone retrieval protocol whereby the vessel, in our case a centre console powerboat, is faced into the wind and the drone reversed toward the bow of the vessel at a height that allows it to be caught by its handles mid-flight, after which the rotors are stopped. This system ensures that the research vessel drifts away from the approaching drone with the wind, rather than into it, keeping more control in the hands of the drone pilot for safe landing.

#### *DNA extraction*

To optimise laboratory processes, a subset of three blow samples were initially chosen to test two different extraction protocols: 1) the same protocol as was used previously (documented in O'Mahony et al. 2024 [1])· whereby the Qiagen

DNeasy Blood and Tissue extraction protocol was slightly modified; and 2) an automated extraction process using the Thermo Fisher Scientific 'KingFisher Duo Prime Purification System'. Based on field metadata, three samples were chosen which were classed as high quality based on droplet size and overall volume in the petri dishes and stemming from individuals which had been sampled in triplicate. The samples were thawed, thoroughly vortexed to homogenise the sample and then divided into two 1000 uL aliquots, from which DNA was extracted using each method. Sample extracts were quantified using the Qubit High-Sensitivity assay. Qiagen-based extracts resulted in higher concentrations, and thus the remaining samples were extracted using this protocol.

For this modified Qiagen extraction protocol, total sample volumes were used (~2000 uL) but initially split across two 1.5 mL Eppendorf tubes to which 180 uL of ATL and 20 uL Proteinase K were added. All samples were incubated at 56°C overnight, rotating slowly. The extraction protocol was modified to use equal volumes of AL and EtOH as present in the sample aliquots. Sample aliquots were pooled in the same spin column, passing 650 uL through the column per round of centrifugation, until the entire sample had been passed through the filter. The remaining standard protocol was followed without modifications, except for two rounds of elution of 75 uL each resulting in a total eluate volume of 150 uL per sample. Blow samples were handled by author ÉOM at the Globe Institute, University of Copenhagen, Denmark.

Tissue samples were extracted using a phenol–chloroform extraction with proteinase K digestion. Between 0.025g and 0.05g of tissue was finely diced and added to a 1.7 mL tube containing 375 uL of proteinase K buffer, 20 uL 10% SDS and 5 uL proteinase K enzyme. Tubes were placed in a rotator and incubated at 54°C for 16-30 hours, turning at low speed. After the incubation period, 400 uL chloroform:isoamyl alcohol was added to each tube, mixed for 2 -5 minutes and the supernatant removed to a new tube. This step was repeated and then 10 uL of 5M NaCl and 600 uL of 95% EtOH was added and held at 20°C overnight. Samples were then spun at maximum speed in a cold centrifuge for 15 minutes to pellet the DNA, then washed twice with 1 mL 70% EtOH, spinning at room temperature for 1-2 minutes. The DNA was let dry for 30 minutes while removing any micro-droplets of EtOH. Once dry, the DNA was resuspended in 50 uL of TE buffer and kept at room temperature overnight before quantification using the spectrophotometer with 2 uL DNA and 98 uL distilled $H_2O$.

***Down sampling of biopsy-sourced genomes***

To explore how depth of coverage affects estimates of heterozygosity and runs of homozygosity, we serially down sampled the biopsy sample raw reads to various target depths using seqtk v.1.4 [82]. We chose to subsample these in steps of 1x and under 5x in steps of 0.25x-0.5x. We did this by calculating the fraction of reads to retain based on the total number of reads available to achieve the target depth of coverage and then used *seqtk sample* on the forward and reverse read FASTQ files for each depth and each biopsy sample. The subsamples were cleaned and mapped as described above.

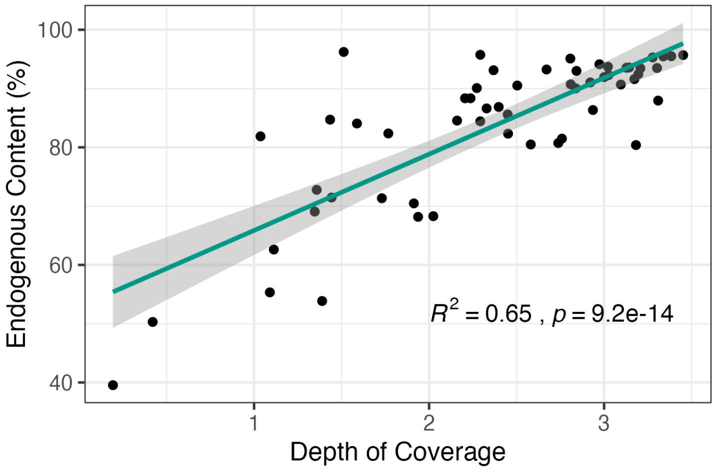


***Figure S1.*** Scatterplot of unmerged blow sample depths of coverage (x-axis) versus each sample's endogenous content (y-axis) with associated Pearson correlation coefficient and p-value, demonstrating a significant positive correlation between the two variables.

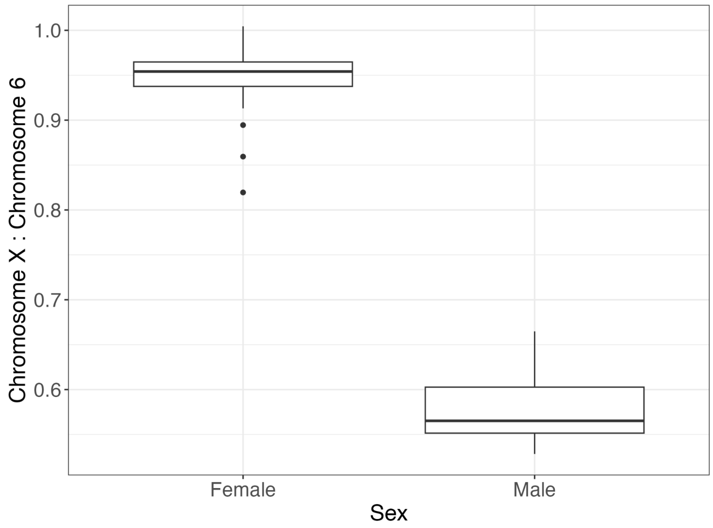


***Figure S2.*** The distribution of sample ratios of number of reads mapping to the X chromosome relative to chromosome six, where a ratio approaching 1 is likely a female (XX sex chromosomes), while a ratio approaching 0.5 will be likely male (XY sex chromosomes).

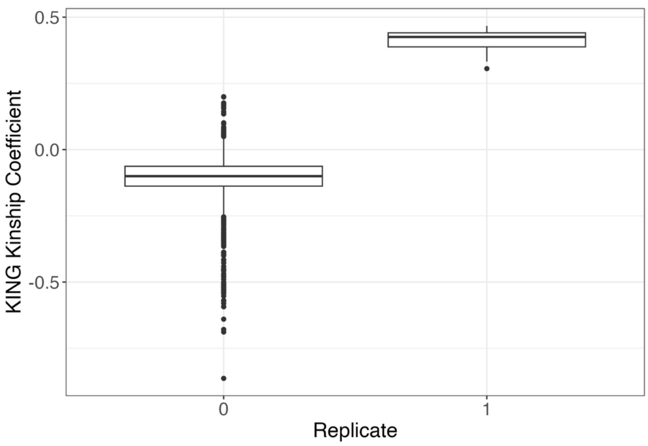


***Figure S3.*** KING-robust kinship coefficients for pairs of blow samples filtered according to resolved false field identifications and sorted according to field replicates (1) and non-replicates (0). This figure represents the post-quality control dataset.

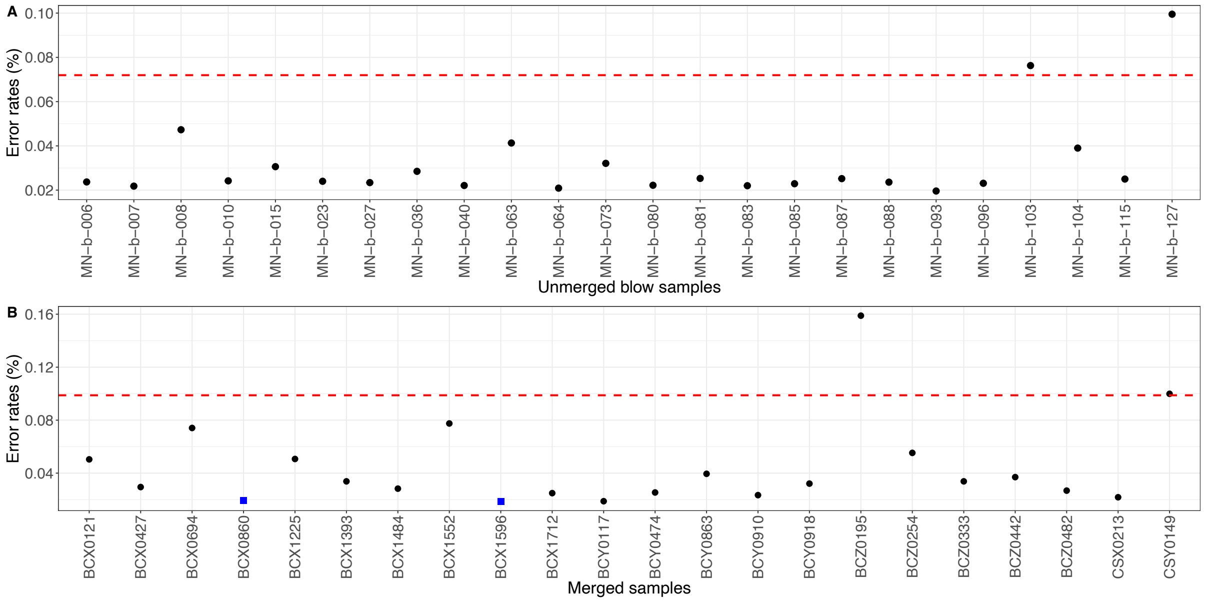


***Figure S4.*** Error rates (%) calculated according to the 'perfect sample approach' in ANGSD. The red dashed lines represent each respective 95th percentile cutoff, above which threshold samples were considered too error prone and discarded from further analysis. **A)** The unmerged blow samples (x-axis) that passed prior QC steps, resulting in the removal of samples MN-b-103 and MN-b-127. **B)** The merged samples (x-axis) that passed prior QC steps, resulting in the removal of BCX0195 and CSY0149. Samples as blue squares are the biopsy-sourced genomes from our study, while the rest are blow sample-sourced.

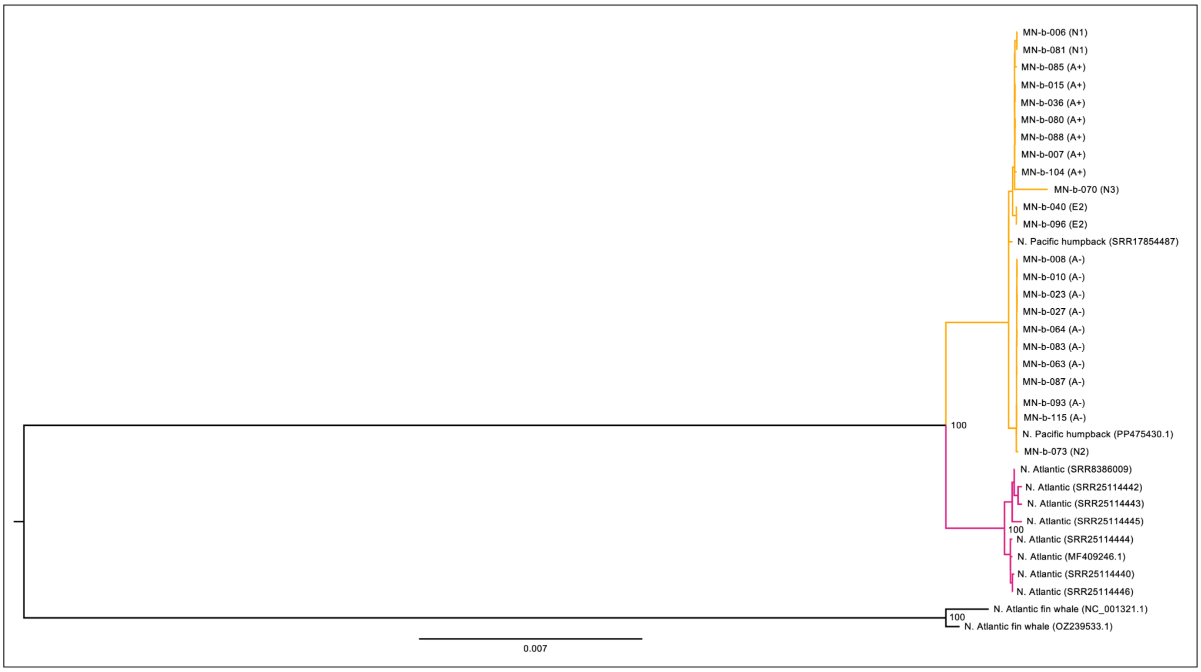


***Figure S5.*** Mitogenome maximum likelihood phylogenetic tree with 100 replicate bootstraps, constructed using the GTR substitution model in IQ-TREE [2]. Samples include the unmerged post-QC blow samples (sample codes MN-b-***) which have their associated control region haplotype noted in brackets, the North Pacific humpback whale reference mitogenome, the DNA Zoo humpback whale reference mitogenome and eight North Atlantic humpback whale mitogenomes (GenBank accession numbers in brackets). Two North Atlantic fin whale mitogenomes were used as an outgroup to root the tree (NC_001321 [3] and OZ239533 (Darwin Tree of Life submission, https://portal.darwintreeoflife.org)).

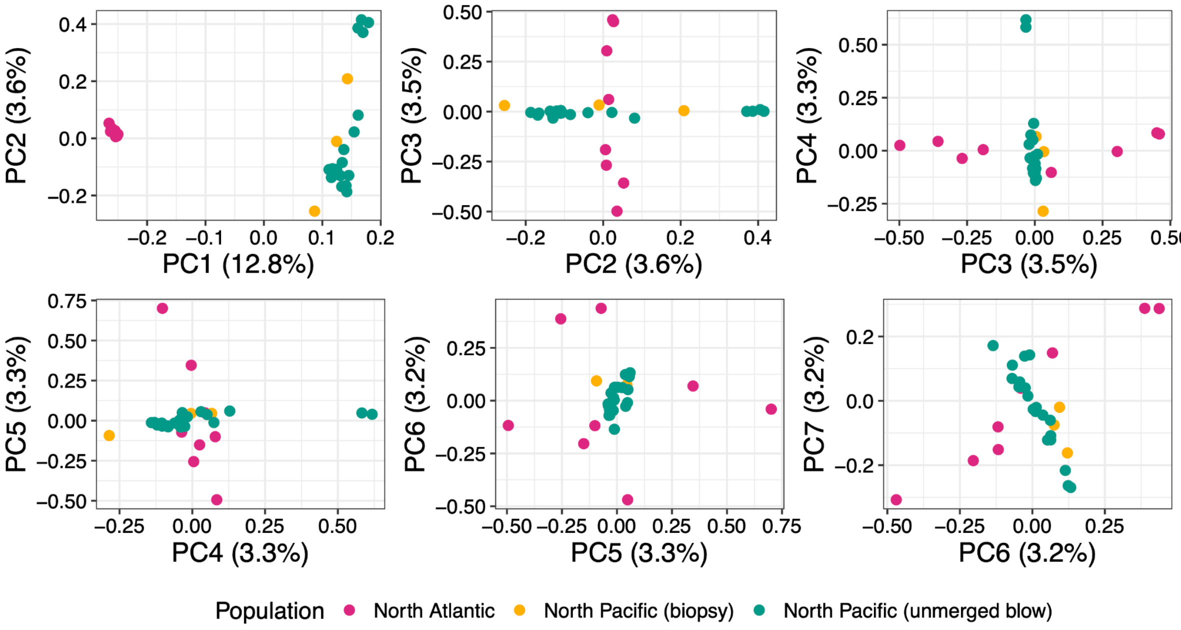


***Figure S6.*** A principal component analysis (PCA) using the unmerged blow samples that passed QC (green) with the biopsy-sourced whole genomes from the North Pacific (yellow) and publicly available whole genomes from the North Atlantic (pink).

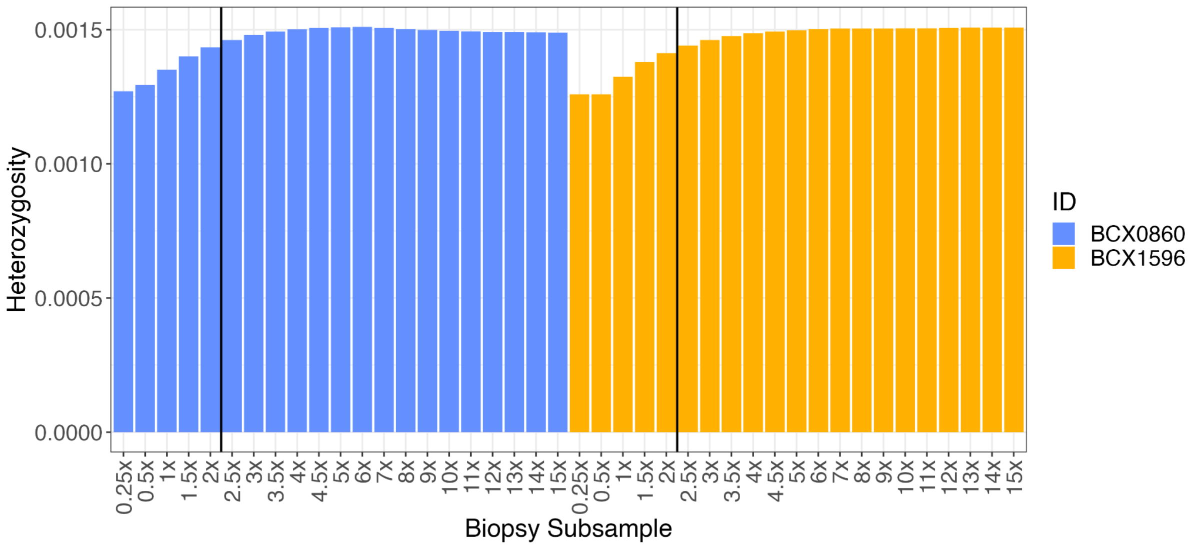


***Figure S7.*** Genome-wide heterozygosity of two humpback whale biopsy samples, subsampled to varying depths of coverage from 15× to 0.25×, demonstrating a decrease in estimated heterozygosity at lower coverages. The black vertical lines mark the mean heterozygosity of the sequenced blow samples analysed in this study.

***Table S1.*** Blow samples stemming from multiple samples of the same individuals. Reported heterozygosity estimates were calculated using a minimum depth threshold of 3×. Whale IDs and blow sample IDs in red with an asterisk were flagged during quality control (QC): either too low depth of coverage, found to be mismatching or fell outside the error rate threshold set by the perfect sample approach and were removed during QC. Depth of coverage (± one standard deviation, 'std. dev.') are reported for unmerged and resolved (i.e. only containing correct IDs) merged samples. Mitochondrial (mtDNA) control region haplotypes are reported according to the haplotypes reported in Baker et al. [4], with novel haplotypes from heteroplasmic sites simply named using the letter N for 'new': 'N1', 'N2' and 'N3'.

| Whale ID | Blow sample IDs | Unmerged depth (± std. dev.) | Unmerged heterozygosity | Resolved merged depth (± std. dev.) | KING kinship value (range) | Resolved merged heterozygosity | mtDNA haplotypes |
|---|---|---|---|---|---|---|---|
| BCX0121 | MN-b-010 | 2.16 | 0.0011 | 3.89 (± 3.36) | 0.36 | 0.0022 | A- |
| | MN-b-109 | 1.73 | 0.0035 | | | | A- |
| BCX0147* | MN-b-006 | 2.58 | 0.0012 | NA | 0.09 | NA | N1 |
| | MN-b-070 | 1.09 | 0.0113 | | | | N3 |
| BCX0171 | MN-b-035 | 3.13 | 0.0012 | 6.00 (± 4.64) | 0.43 | 0.0016 | A- |
| | MN-b-043 | 2.45 | 0.0013 | | | | A- |
| BCX0427 | MN-b-002 | 3.31 | 0.0016 | 6.33 (± 4.64) | (-0.13 to 0.44) | 0.0016 | A+ |
| | MN-b-089* | 1.36 | 0.0040 | | | | E2 |
| | MN-b-090 | 3.02 | 0.0013 | | | | A+ |
| BCX0694 | MN-b-013* | 0.22 | NA | 2.17 (± 2.36) | NA | NA | N1 |
| | MN-b-081 | 1.96 | 0.0013 | | | | N1 |
| BCX0860 | MN-b-027 | 2.97 | 0.0016 | 6.14 (± 4.36) | 0.42 | 0.0016 | A- |
| | Biopsy (3× subsample) | 15.06 | 0.0015 | | | | A- |
| BCX1225 | MN-b-045 | 2.27 | 0.0013 | 4.78 (± 3.65) | 0.39 | 0.0022 | A- |
| | MN-b-103 | 2.50 | 0.0031 | | | | A- |
| BCX1393 | MN-b-018 | 2.76 | 0.0021 | 6.14 (± 4.50) | 0.43 | 0.0017 | E2 |
| | MN-b-096 | 3.38 | 0.0014 | | | | E2 |

| | | | | | | | |
|---|---|---|---|---|---|---|---|
| BCX1484 | MN-b-004 | 1.39 | 0.0018 | 10.96 (± 7.45) | (0.44 to 0.44) | 0.0017 | A- |
| | MN-b-023 | 3.21 | 0.0014 | | | | A- |
| | MN-b-079 | 3.17 | 0.0012 | | | | A- |
| | MN-b-095 | 3.20 | 0.0013 | | | | A- |
| BCX1552 | MN-b-020 | 1.77 | 0.0021 | 4.06 (± 3.42) | 0.38 | 0.0031 | A- |
| | MN-b-127 | 2.29 | 0.0039 | | | | A- |
| BCX1596 | MN-b-066 | 2.29 | 0.0014 | 5.10 (± 3.93) | (0.43 to 0.47) | 0.0015 | A- |
| | MN-b-087 | 2.81 | 0.0014 | | | | A- |
| | Biopsy | 15.85 | 0.0015 | NA | | NA | A- |
| BCX1712 | MN-b-007 | 2.94 | 0.0012 | 5.61 (± 4.27) | 0.43 | 0.0014 | A+ |
| | MN-b-078 | 2.67 | 0.0015 | | | | A+ |
| BCX2052 | MN-b-015 | 3.02 | 0.0016 | 5.35 (± 4.21) | 0.39 | 0.0022 | A+ |
| | MN-b-082 | 2.33 | 0.0029 | | | | A+ |
| BCY0117 | MN-b-009 | 2.74 | 0.0011 | 6.02 (± 4.46) | 0.44 | 0.0013 | A- |
| | MN-b-093 | 3.28 | 0.0013 | | | | A- |
| BCY0474 | MN-b-001* | 2.02 | 0.0022 | 4.27 (± 3.81) | (0.26 to 0.38) | 0.0014 | A- |
| | MN-b-084 | 1.43 | 0.0014 | | | | A+ |
| | MN-b-085 | 2.84 | 0.0013 | | | | A+ |
| BCY0863 | MN-b-048 | 1.11 | 0.0017 | 2.70 (± 2.56) | 0.36 | 0.0019 | A+ |
| | MN-b-104 | 1.59 | 0.0019 | | | | A+ |
| BCY0910 | MN-b-055 | 3.30 | 0.0013 | 6.64 (± 4.67) | 0.45 | 0.0015 | A+ |
| | MN-b-088 | 3.34 | 0.0014 | | | | A+ |
| BCY0918 | MN-b-036 | 2.24 | 0.0015 | 3.27 (± 3.08) | 0.34 | 0.0017 | A+ |

| | | | | | | | |
|---|---|---|---|---|---|---|---|
| | MN-b-092 | 1.04 | 0.0018 | | | | A+ |
| BCY0965 | MN-b-003 | 3.18 | 0.0012 | 8.72 (± 6.41) | (0.42 to 0.44) | 0.0018 | A- |
| | MN-b-083 | 3.14 | 0.0013 | | | | A- |
| | MN-b-099 | 2.40 | 0.0029 | | | | A- |
| BCZ0195* | MN-b-008 | 1.94 | 0.0018 | 3.28 (± 3.04) | 0.31 | 0.0058 | A- |
| | MN-b-091 | 1.35 | 0.0110 | | | | A- |
| BCZ0254 | MN-b-040 | 3.10 | 0.0013 | 4.54 (± 3.60) | 0.36 | 0.0024 | E2 |
| | MN-b-069 | 1.44 | 0.0049 | | | | E2 |
| BCZ0333 | MN-b-111 | 2.81 | 0.0020 | 5.81 (± 4.38) | 0.43 | 0.0017 | A- |
| | MN-b-115 | 3.00 | 0.0014 | | | | A- |
| BCZ0442 | MN-b-063 | 2.37 | 0.0019 | 3.88 (± 3.10) | 0.40 | 0.0018 | A- |
| | MN-b-106 | 1.51 | 0.0015 | | | | A- |
| BCZ0482 | MN-b-064 | 3.45 | 0.0013 | 5.66 (± 4.22) | 0.43 | 0.0015 | A- |
| | MN-b-126 | 2.21 | 0.0018 | | | | A- |
| CSX0213 | MN-b-028 | 2.84 | 0.0013 | 5.76 (± 4.26) | 0.43 | 0.0014 | A+ |
| | MN-b-080 | 2.92 | 0.0013 | | | | A+ |
| CSY0149* | MN-b-073 | 2.45 | 0.0016 | 4.36 (± 3.50) | 0.34 | 0.0039 | N2 |
| | MN-b-074 | 1.91 | 0.0071 | | | | N2 |

***Table S2.*** Potential occurrence of heteroplasmy in samples differing from known North Pacific mtDNA control region D-loop haplotypes. Even samples of low mitogenome coverage (MN-b-013 and MN-b-117) show heteroplasmy at the correct site and are in agreement with their higher coverage replicate samples. Samples with an asterisk were later determined to have a false field-ID, due to the discrepancies in heteroplasmic sites and KING-robust kinship coefficients.

| **Sample** | **ID** | **Haplo-types** | **Coverage (mtDNA)** | **Pos 15,640 Ref: T** | **Depth 15,640** | **Pos: 15,802 Ref: C** | **Depth 15,802** | **Pos: 15,822 Ref: A** | **Depth 15,822** |
|---|---|---|---|---|---|---|---|---|---|
| MN-b-006* | BCX0147* | N1 | 119.7x | C:T 86:33 | 119x | C | 101x | A | 101x |
| MN-b-013 | BCX0694 | N1 | 8.3x | C:T 5:1 | 6x | C | 2x | A | 3x |
| MN-b-081 | BCX0694 | N1 | 94.1x | C:T 51:22 | 73x | C | 58x | A | 68x |
| MN-b-073 | CSY0149 | N2 | 129.4x | T | 92x | T:C 47:47 | 94x | A | 99x |
| MN-b-074 | CSY0149 | N2 | 103.7 | T | 78x | T:C 35:34 | 69x | A | 74x |
| MN-b-117 | CSY0149 | N2 | 9.07x | T | 5x | T:C 2:1 | 3x | A | 2x |
| MN-b-070* | BCX0147* | N3 | 63.9x | T | 49x | C | 49x | G:A 25:23 | 48x |

***Table S3.*** Pairs of samples flagged as highly related in the NgsRelate run (KING-robust kinship coefficient > 0.125) using the merged blow samples, biopsies and publicly available data. Samples of higher coverage were retained for downstream analyses (and those removed highlighted with an asterisk).

| **ID1 (depth ± SD)** | **ID2 (depth ± SD)** | **KING** |
|---|---|---|
| BCX1484 (11.0× ± 7.5) | BCY0965 * (8.7× ± 6.4) | 0.135 |
| BCX0171 * (6.0× ± 4.6) | BCY0117 (6.0× ± 4.5) | 0.213 |
| BCX0427 (6.3× ± 4.6) | BCX2052 * (5.4× ± 4.2) | 0.214 |
| BCX0860 merged blow with 3× subsampled biopsy * (6.1× ± 4.4) | BCX0860 biopsy sample (15.1× ± 8.0) | 0.461 |
| BCX1596 merged blow samples * (5.1× ± 3.9) | BCX1596 biopsy sample (15.8× ± 8.5) | 0.482 |

***Table S4.*** Publicly available nuclear and mitochondrial genome resequencing data used for ground truthing the blow sample generated whole genomes in this study. Samples chosen from Suárez-Menéndez et al. [5] were from the parental generation, to avoid related dyads.

| **Accession number** | **BioProject** | **Ocean basin** | **Depth (± var)** | **Heterozygosity** | **Citation** |
|---|---|---|---|---|---|
| SRR17854487 | PRJNA512907 | Pacific | 61.8× (± 21.8) | 0.0014 | DNA Zoo [6] |
| SRR8386009 | PRJNA509641 | North Atlantic | 8.2× (± 6.0) | 0.0022 | Tollis et al. [7] |
| SRR5665639 | PRJNA389516 | North Atlantic | 18.8× (± 10.3) | 0.0015 | Árnason et al. [8] |
| SRR25114440 | PRJNA990679 | North Atlantic | 25.6× (± 11.6) | 0.0014 | Suárez-Menéndez et al. [5] |
| SRR25114442 | PRJNA990679 | North Atlantic | 26.7× (± 11.6) | 0.0014 | Suárez-Menéndez et al. [5] |
| SRR25114443 | PRJNA990679 | North Atlantic | 26.7× (± 11.8) | 0.0015 | Suárez-Menéndez et al. [5] |
| SRR25114444 | PRJNA990679 | North Atlantic | 26.4× (± 11.6) | 0.0014 | Suárez-Menéndez et al. [5] |
| SRR25114445 | PRJNA990679 | North Atlantic | 26.0× (± 11.4) | 0.0014 | Suárez-Menéndez et al. [5] |
| SRR25114446 | PRJNA990679 | North Atlantic | 26.7× (± 11.5) | 0.0014 | Suárez-Menéndez et al. [5] |
